# Efficient Identification of Phylogenetically Informative Alignment Sites via Sparse Learning

**DOI:** 10.1101/2025.07.24.666198

**Authors:** Carlos G. Schrago

## Abstract

Identifying phylogenetically informative sites in multiple sequence alignments is critical for accurate tree reconstruction and efficient data curation in phylogenomics. Existing approaches that measure phylogenetic information often rely on predefined topologies or heuristic criteria, limiting their generality and interpretability. Here, we introduce a novel, topology-agnostic framework for quantifying site-wise phylogenetic information using sparse learning via Lasso (Least Absolute Shrinkage and Selection Operator) regression. By modeling site log-likelihoods as predictors of the tree likelihood across a large ensemble of random topologies, our approach isolates the minimal subset of sites that meaningfully contribute to phylogenetic signal. We validate the method using both simulated and empirical mammalian datasets, demonstrating that Lasso-selected sites yield topologies nearly identical to those inferred from full alignments. For computational efficiency, we show that a simple entropy-based proxy (Shannon *H* ≥ 0.5) approximates Lasso results with high fidelity, enabling rapid site-level assessments. Importantly, our definition of phylogenetically informative sites provides an objective metric that can serve as a gold standard to evaluate commonly used alignment filtering tools. These findings establish sparse learning as a principled, scalable, and practical approach for assessing and optimizing phylogenetic data.

## Introduction

Phylogenies are essential for addressing most evolutionary questions, and the development of phylogenetic methods has steadily moved toward a solid statistical foundation (Felsenstein, 2004; Yang, 2014). A central challenge in this framework is to quantify the phylogenetic information content of multiple sequence alignments (MSAs)—that is, its capacity to resolve evolutionary parameters such as tree topology or branch lengths with statistical confidence. Following the foundational work of Felsenstein (Felsenstein, 1981), phylogenetic inference has relied heavily on the maximum likelihood framework, where information is often conceptualized in terms of Fisher information (Edwards, 1972). This perspective has underpinned developments ranging from model selection (Felsenstein, 1988; Goldman, 1993) to hypothesis testing using likelihood ratio and topology comparison tests (Kishino and Hasegawa, 1989; Shimoidara, 2002).

Yet, despite these advances, a common practical challenge frequently encountered in large-scale phylogenomic studies is identifying which sites of an MSA or which loci significantly contribute to phylogenetic resolution. This question differs fundamentally from classical model selection, which is typically implemented in phylogenetics and usually evaluates model fit (mostly substitution models) for a single, complete alignment (Goldman, 1993). To address the challenges of site selection, various approaches have been proposed. For instance, a measure of phylogenetic informativeness profile of a locus was proposed to quantify the capacity of an MSA to resolve tree branches over several evolutionary timescales (Townsend, 2007). More recently, gene-wise or site-wise log-likelihood score comparisons calculate how much better a locus or site fits one candidate topology over another, often using log-likelihood difference values based on pre-specified tree topologies (Shen *et al*., 2017; Walker *et al*., 2018). These approaches require a priori definition of competing topologies and, thus, cannot be employed as general tools to quantify the intrinsic phylogenetic information of a locus or alignment site. Few methods provide objective metrics for assessing gene or site informativeness without relying on a limited set of predefined topologies or hypotheses (Criscuolo and Gribaldo, 2010; Dress *et al*., 2008).

Recently, Haag et al. (Haag *et al*., 2022) trained a machine learning model that is topology-agnostic and is capable of evaluating the difficulty of resolving a maximum likelihood tree topology based on a full MSA. At the site level, however, the assessment of phylogenetic information content at individual sites often depends on assumptions that warrant more rigorous statistical examination. As a result, an explicit evaluation of the performance of approaches that use site-wise calculations, such as the gap frequency or entropy of alignment sites, is complicated. These approaches are generally employed for filtering alignments in phylogenomic pipelines to ensure faster computational times by identifying poorly aligned or weakly informative regions in analytical workflows (Steenwyk *et al*., 2020; Talavera and Castresana, 2007).

To address this limitation, we introduce a framework for assessing site-wise phylogenetic information based on sparse learning via the Lasso (Least Absolute Shrinkage and Selection Operator) regressor (Tibshirani, 1996). Sparse learning methods penalize regression coefficients to encourage sparsity, effectively selecting a minimal subset of parameters that explain the response variable (Hastie *et al*., 2016). Lasso has recently been employed to address several problems in phylogenetics (Ecker *et al*., 2022; Kumar and Sharma, 2021; Sharma and Kumar, 2024), and its models are generally more interpretable than parameter-rich deep learning models.

We implement Lasso to model site-wise log-likelihoods as predictors of the tree log-likelihood, enabling the identification of the subset of informative sites. This approach leverages Lasso’s sparsity-inducing nature to isolate the sites most critical for resolving phylogenetic relationships. We also suggest that the number of Lasso-selected sites can be used to quantify the phylogenetically effective alignment length. This approach provides a principled, model-aware metric of site informativeness with practical utility for alignment trimming and marker selection in phylogenetics.

## Materials and Methods

Our central idea is that sparse learning via Lasso regression can objectively identify a minimal subset of alignment sites that effectively capture phylogenetic information without relying on predefined topologies. We implemented our approach as follows.

### Phylogenetic information content of sequence sites using Lasso

We borrowed the idea from Ecker et al. (Ecker *et al*., 2022) to identify alignment sites that contribute to the tree likelihood using Lasso regression (Tibshirani, 1996). In this framework, site-specific log-likelihoods are treated as predictors (features), while the overall tree log-likelihood serves as the response variable. As is standard in phylogenetic analyses, sites are assumed to be independent and identically distributed. Let *p* denote the number of nucleotide sites in the alignment. Thus, a linear regression model can be expressed as In 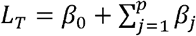 In *L*_*j*_. Where In *L*_*T*_ is the tree log-likelihood, In *L*_*j*_ is the log-likelihood of the *j*^th^ site, *β*_*j*_ is the corresponding regression coefficient, and *β*_0_ is the intercept.

For each alignment, to generate observations and estimate the coefficients of the linear model, we produced 10,000 random trees (including both topology and branch lengths) using IQ-TREE2 (Minh *et al*., 2020), employing the ‘-r’ flag. For each random tree, the tree likelihood was calculated using the ‘--sitelh’ command to record the log-likelihoods of each alignment site. The Lasso regression framework enables the estimation of regression coefficients (*β*) that may be precisely zero. Consequently, alignment sites with non-zero *β* coefficients are interpreted as having a meaningful impact on the response variable, the tree log-likelihood. Lasso analyses were performed in R using the *glmnet* package. The site-wise coefficients were inferred by minimizing the following objective function.

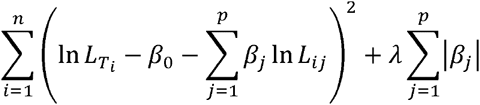

Here, *n* denotes the number of observations (tree topology replicates), In 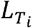 the log-likelihood of the *i*^th^ tree, *p* is the number of alignment sites, *β*_*j*_ is the coefficient associated with the *j*^th^ site, and *L*_*ij*_ the log-likelihood of the *j*^th^ site of the *i*^th^ tree. The first term of the above formula represents the residual sum of squares (RSS), i.e., the sum of squared differences between the observed tree log-likelihoods 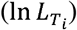 and the predicted values from the linear model 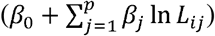,. The second term, 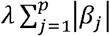, corresponds to the L1-norm of the regression coefficients and incorporates the tuning parameter *λ*, which controls the degree of penalization applied to the model. When *λ* = 0, the Lasso model reduces to ordinary least squares; as *λ* increases, more coefficients are shrunk towards zero, promoting sparsity. The optimal *λ* for each alignment was determined through cross-validation (James *et al*., 2017). An alignment site was classified as uninformative if its associated *β* coefficient was zero, and informative otherwise.

### Simulated and empirical datasets

We simulated 1,000 alignments under the GTR+G4 model with indels using the AliSim sequence simulator implemented in IQ-TREE2 (Ly-Trong *et al*., 2022). To closely approximate real biological sequences, parameters for the GTR model, alignment lengths, and the alpha parameter of the Gamma distribution were derived from empirical estimates of mammalian coding sequences available in the OrthoMam v12 database (Allio *et al*., 2024). Additionally, we calculated the frequencies of each indel size in OrthoMam sequences and used a Zipf distribution to generate realistic gap-containing alignments. All simulations were based on empirical topologies also obtained from OrthoMam, which broadly reflect the diversification of mammalian orders and families. The average alignment length was 1,986.3 bp, ranging from 357 to 13,383. To evaluate the effect of taxon sampling on the performance of our approach, we generated two datasets by subsampling sequences from the simulated alignments: one with 20 taxa (1,000 alignments) and another with 100 taxa (250 alignments). Each alignment was analyzed using the Lasso procedure described above.

For empirical analyses, we retrieved 17,439 orthologous gene alignments representing approximately 480 mammalian species, assembled using the TOGA pipeline (Kirilenko *et al*., 2023). From this dataset, we randomly selected 1,000 gene alignments and, for each, randomly sampled 20 taxa for downstream analysis. Although both OrthoMam and TOGA provide 1-to-1 ortholog alignments, we opted to use TOGA for empirical evaluation due to its broader taxonomic coverage (480 species vs. 190 in OrthoMam).

### Evaluating the performance of Lasso

The performance of Lasso was evaluated in simulated and empirical datasets, as described below. In both cases, we ran the cross-validation procedure for estimating the optimal *λ* parameter twice to ensure the repeatability of our approach. We then measured repeatability as the proportion of sites consistently classified into the same category (informative or uninformative) across the two independent runs. Given that the substitution model influences the site log-likelihoods, we examined the robustness of our approach to model misspecification. To that end, we conducted Lasso analyses on simulated data using the simplest substitution model, JC69, which was not used to generate those alignments. Robustness was quantified as the proportion of sites receiving the same classification under both the correct and incorrect substitution models.

We assembled two simulated datasets, each comprising 20 and 100 terminals. This allowed us to investigate whether Lasso-selected sites affected the accuracy of phylogenetic inference. The Robinson-Foulds distance (Robinson and Foulds, 1981) was calculated between the true topology and the IQ-TREE2-generated maximum likelihood topologies inferred from (*i*) the complete alignment, as well as the alignments containing only (*ii*) informative or (*iii*) uninformative sites. Since the simulated alignments were derived from empirical trees with more than 100 terminals, the true topologies were obtained by pruning these full trees to match the taxa present in the 20- and 100-taxa datasets. In addition to RF distances, we calculated the proportion of significantly resolved branches (*P* > 0.95) by the parametric aLRT statistic (Anisimova and Gascuel, 2006) for all inferred trees in both simulated and empirical datasets.

### Approximating Lasso results with a simple metric

Because the Lasso procedure can be both time and computationally intensive, we explored whether the site-wise Shannon entropy (*H*), which is fast and easy to compute, could serve as a proxy for the Lasso-based site classification. Treating indels as a fifth character state, entropy was calculated for each nucleotide alignment column as 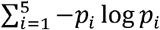, where *p*_*i*_ denotes the frequency of the *i*^th^ character in an alignment column. To determine an optimal entropy threshold (cut-off) for classifying sites as informative or not, we identified the value of *H* that maximized the agreement between entropy-based classification and the Lasso-derived results.

To achieve this, we employed a binary coding for the Lasso result, assigning 1 to informative sites and 0 to uninformative sites. We then fitted a logistic regression model using the *glm* function of R, with the site-wise entropy as the predictor and the Lasso binary-coded classification as the response. The threshold site-wise entropy value was determined by calculating the ratio −*a/b*, where *a* is the intercept and *b* the slope of the logistic regression model, respectively. This value corresponds to the entropy at which the model predicts a 50% probability of being classified as informative.

### Using site-wise phylogenetic information for comparing alignments and dataset filtering

Lasso results can be leveraged to compare the phylogenetic information content across different genes. Specifically, the number of sites selected as informative by Lasso provides an estimate of the effective alignment length, serving as a measure for the information content of a multiple sequence alignment (MSA). In phylogenomics pipelines, this metric can guide the selection of phylogenetically informative loci. To assess the relevance of this measure, we verified whether the number of informative sites inferred by Lasso correlates with the degree of phylogenetic difficulty estimated by the Pythia predictor (Haag *et al*., 2022). This predictor yields a continuous score ranging from 0 (indicating high phylogenetic resolution) to 1 (indicating alignments likely to produce poorly resolved trees). Accordingly, we expected a negative correlation between the number of informative sites and Pythia scores.

The results of the Lasso regressor can also be applied to both filter alignments and evaluate alignment trimming tools. Following this rationale, we filtered our alignments to retain only sites that informed the tree likelihood as identified by Lasso. Using these alignments containing only informative sites as the standard, we compared them with those generated by TrimAl (Capella-Gutiérrez *et al*., 2009) and BMGE (Criscuolo and Gribaldo, 2010), both under default settings. These trimming tools were selected because they implement alignment filtering under distinct principles. BMGE selects alignment columns by computing an entropy-like score weighted with biological substitution matrices to identify regions with unexpectedly high variability, while trimAl applies thresholds based on gap presence, similarity, and consistency scores. To quantify the agreement between Lasso and the other methods, we calculated the proportion of sites classified identically— i.e., whether a site was retained (informative) or removed (uninformative) in both filtered alignments.

## Results

In simulated datasets, the average proportion of sites classified as informative by Lasso was 15.1% (95% quantiles: 6.4%–29.2%) for the 20-taxa dataset and 24.6% (9.4%–39.8%) for the 100-taxa dataset, indicating that increasing the number of taxa led to a higher frequency of informative sites. In the empirical dataset, the average proportion of informative sites was 22.5% (9.4%–41.0%). Across all datasets, the number of informative sites was positively correlated with the total alignment length, with *R*^2^ values near or exceeding 0.6 and *p* < 0.001 (**Figure 1**). Thus, sampling more sites increased the likelihood of capturing phylogenetically informative positions.

**Figure 1.**
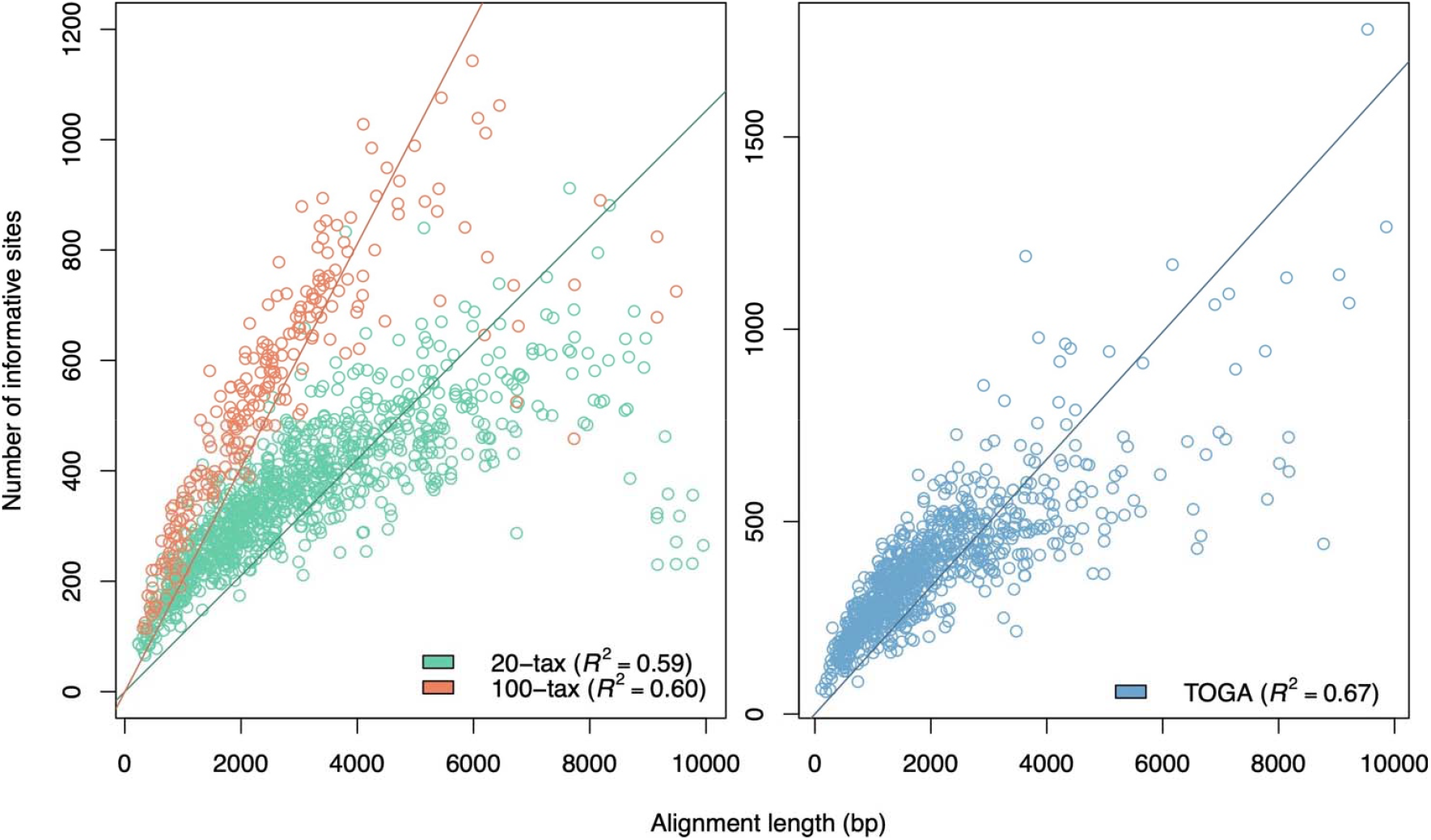
Correlation between the number of Lasso-selected informative sites and alignment length across all datasets (*p* < 0.01). Regression lines were fitted using the robust linear model function (*rlm*) to control for the effects of outliers.

The average site-wise agreement between the two independent cross-validation runs was 98.3% (range: 96.5%–99.5%), indicating a high level of consistency in site classification. When analyses were conducted using an incorrect nucleotide substitution model (JC69), which poorly fit these alignments, the average site-wise agreement between classifications under the correct and incorrect models was 85.2% (78.4%–90.1%). This result suggests that the Lasso regressor is reasonably robust to model misspecification. Finally, the average agreement in site classification between the 20- and 100-taxa datasets was 85.5% (77.8%–91.1%), reflecting the influence of taxon sampling on Lasso-based site selection.

The RF topological distances to the true tree were similar for trees inferred from complete alignments and those based solely on informative sites, regardless of the number of taxa. For the 20- and 100-taxa datasets, the average RF distances between the inferred and true topologies were 4.2 and 26.5, respectively, when using complete alignments, and 5.1 and 28.3 when using only Lasso-selected informative sites. In contrast, trees reconstructed from sites classified as uninformative by Lasso showed substantially higher RF distances—averaging 11.3 for the 20-taxa dataset and 101.0 for the 100-taxa dataset—indicating a marked reduction in phylogenetic accuracy (**Figure 2a**).

**Figure 2.**
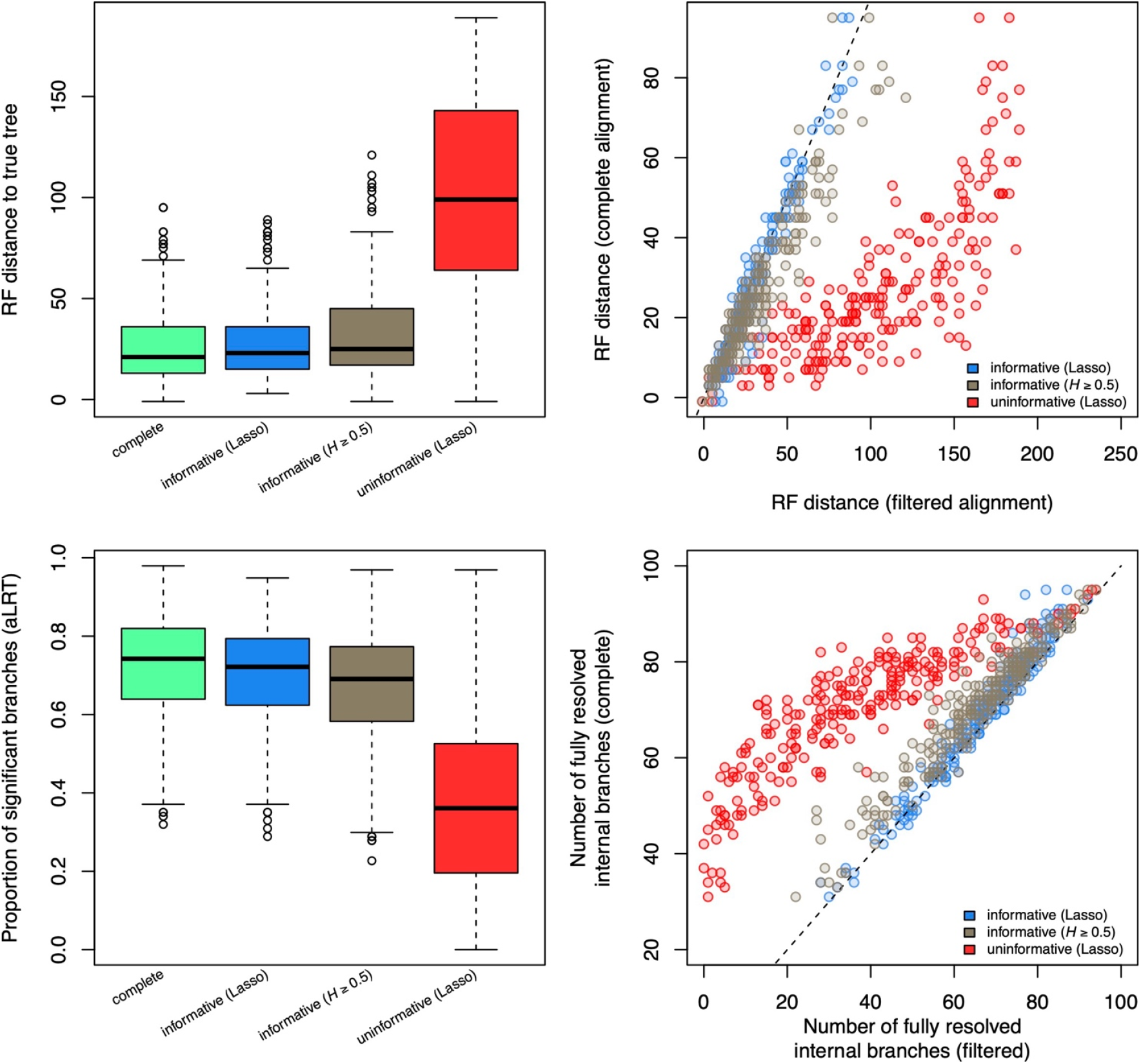
Performance of filtered alignments (Lasso and Shannon entropy) compared to complete alignments in recovering the true topology. Results shown are for the 100-taxa dataset only. (a) Boxplots of Robinson–Foulds (RF) distances between inferred trees and the true tree for different site compositions. (b) Correlation between RF distances of complete and filtered alignments composed of Lasso-selected sites or sites with *H*□≥□0.5 (*R*^2^ > 0.9, *p* < 0.01). The dashed line represents the *y* = *x* identity line. (c) Proportion of branches with parametric aLRT ≥ 0.95 (Anisimova and Gascuel, 2006) in topologies inferred from each alignment type. (d) Correlation between the number of fully resolved branches (aLRT ≥ 0.95) between the tree inferred by complete alignment and those inferred with filtered alignments. Topologies inferred from Lasso-selected sites and sites with *H* ≥ 0.5 are highly correlated with those inferred from complete alignments (*R*^2^ > 0.9, *p* < 0.01). The dashed line indicates the *y* = *x* relationship.

In both simulated and empirical datasets, trees inferred from Lasso-selected informative sites exhibited proportions of branches with aLRT support values greater than 0.95 that were comparable to those observed in trees reconstructed from complete alignments. In contrast, the proportion of fully resolved nodes declined markedly when trees were inferred using only sites classified as uninformative by Lasso (**Figure 2b**). On a case-by-case basis, the number of well-supported nodes in trees derived from informative-site alignments closely matched those from the full alignments. By comparison, alignments composed exclusively of uninformative sites consistently performed worse across all evaluated metrics. (**Figure 2c**).

The mean Shannon entropy of sites selected by Lasso was similar for all datasets investigated: 0.71 for the 20-taxa, 0.64 for the 100-taxa, and 0.65 for the TOGA empirical dataset. In contrast, sites classified as uninformative tended to have lower entropy values, with distributions skewed toward minimal variation (**Figure 3a**). Based on the 100-taxa dataset, the entropy threshold that best separated informative from uninformative sites according to Lasso was *H =* 0.54, which was rounded to 0.5 for simplicity. This was a conservative cut-off, because both 20-taxa and empirical datasets exhibited thresholds greater than 0.5 (**Figure 3b**). Using this operational cut-off, sites were considered as informative if they exhibited *H* ≥ 0.5. This simple rule resulted in a high degree of concordance with the Lasso-based classification, yielding 88.3% agreement for the 100-taxa dataset, 86.3% for the 20-taxa dataset, and 83.1% for the TOGA empirical dataset.

**Figure 3.**
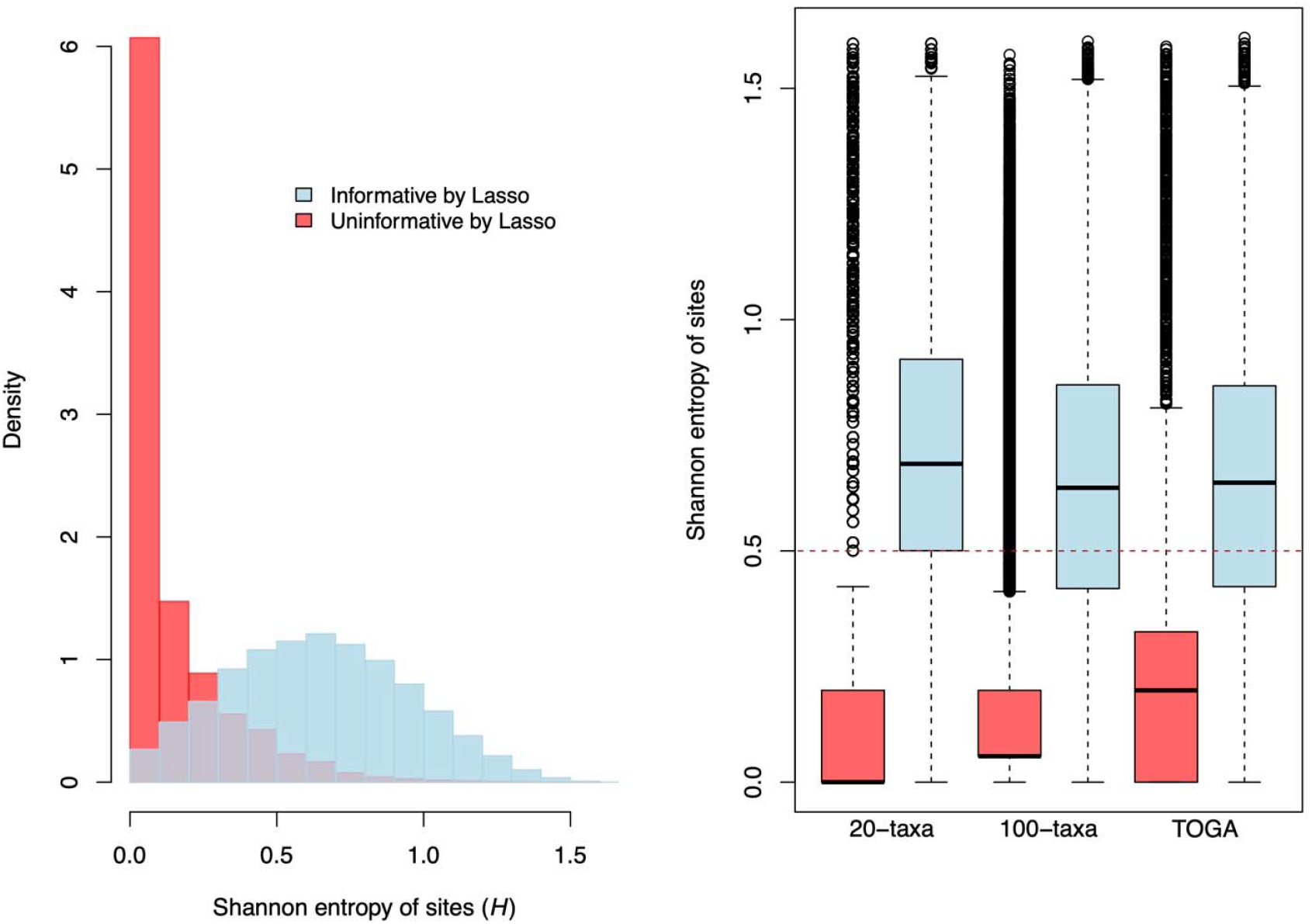
Shannon entropy distributions for sites classified by Lasso as informative or uninformative. Results shown are for the 100-taxa dataset only. (a) Histogram of entropy values. (b) Boxplots of entropy values across all datasets. The red dashed line marks the operational cut-off of *H*□=□0.5.

Consistent with the results obtained using Lasso-selected informative sites, RF distances between the true trees and those inferred from alignments composed of sites with *H* ≥ 0.5 were also close to 50 on average (**Figure 2a**). Moreover, RF distances derived from Lasso-based informative alignments were highly correlated with those from the *H*□≥□0.5 alignments (*R*^2^ = 0.96, *p* < 0.01). In contrast, alignments composed of *H <* 0.5 sites yielded trees with an average RF distance of 144.2, slightly lower than the 160.9 observed for alignments containing sites classified as uninformative by Lasso. This was expected, because the *H <* 0.5 set included a small proportion of sites that Lasso identified as informative (**Figure 3**). Notably, RF distances from the *H*□<□0.5 and Lasso-uninformative alignments were also strongly correlated (*R*^2^ = 0.93, *p* < 0.01).

Among the sites classified as informative by Lasso in the 100-taxa dataset, 86.6% and 83.2% were also retained by trimAl and BMGE, respectively. Both trimAl and BMGE produced substantially longer alignments than those filtered using Lasso. On average, trimAl and BMGE retained 55.0% and 52.3% of the original sites, respectively, while the Lasso approach retained 24.6%. Alignments filtered using the Shannon entropy cut-off as a proxy for Lasso were the shortest, with an average retention of 18.2% (**Figure 4a**). Despite their reduced length, phylogenies inferred from Lasso- and entropy-filtered alignments exhibited topological distances to the true tree that were comparable to those obtained using trimAl and BMGE (**Figure 4b**). This indicates a higher phylogenetic efficiency for the Lasso-based approaches, i.e., for a given filtered alignment length, they yielded more accurate trees (**Figure 4c**). In most cases, RF distances between trees inferred from Lasso-filtered alignments and the true tree were less than or equal to those obtained using trimAl or BMGE (**Table S1**).

**Figure 4.**
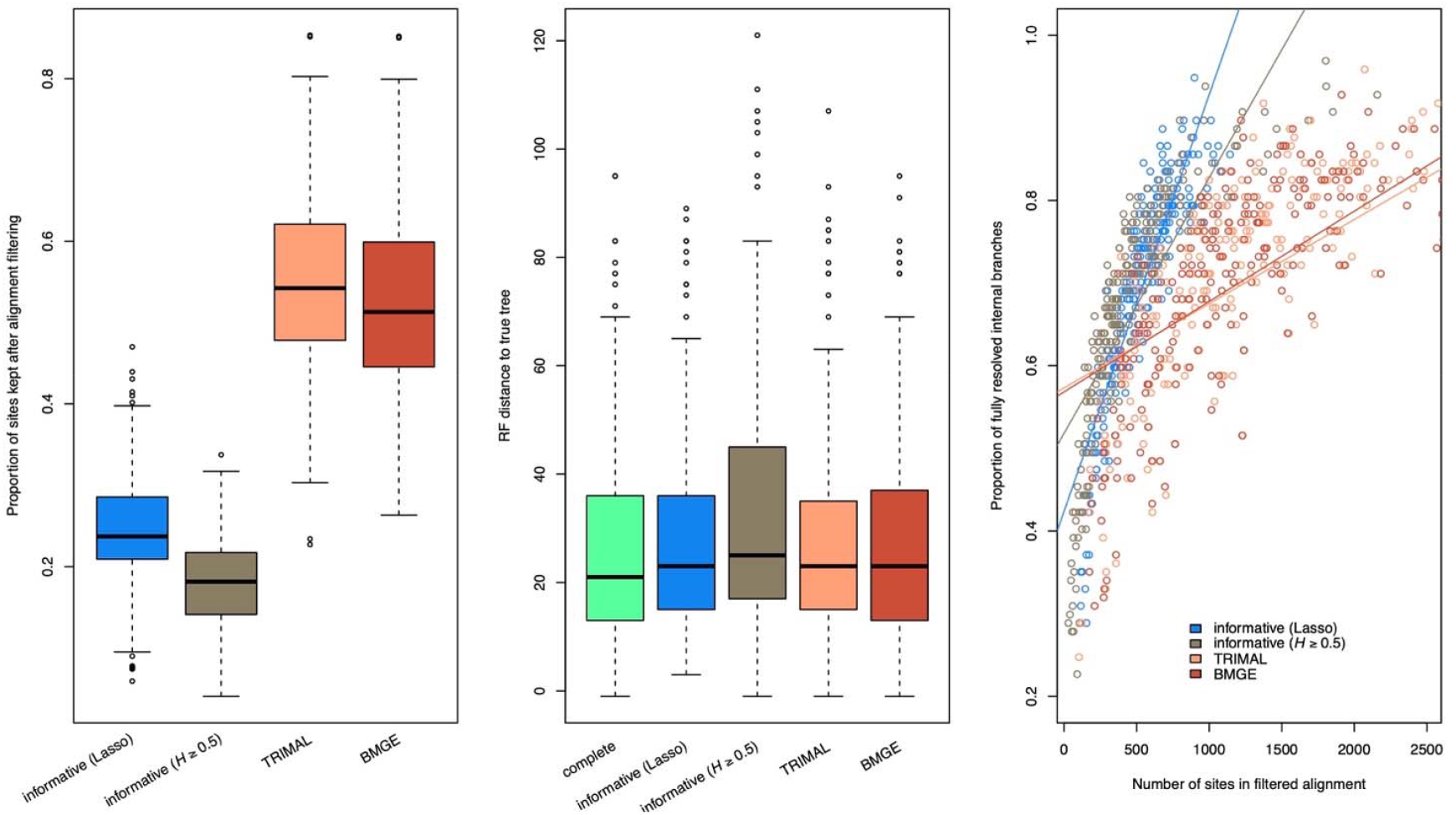
Comparison of Lasso-informed alignment filtering versus Trimal and BMGE. Results shown are for the 100-taxa dataset only. (a) Proportion of sites retained by each method. (b) Boxplots of the Robinson-Foulds (RF) distance between the true tree topology used to simulate alignments and for each alignment site composition. (c) Correlation between the proportion of fully resolved branches (aLRT ≥ 0.95) and the number of sites in filtered alignments by each method. Despite shorter alignments, Lasso-based filtering produced a higher proportion of well-supported branches.

We observed an inverse correlation between the number of informative sites and the difficulty of resolving the tree topology, as estimated by the Pythia predictor (**Figure 5**). Notably, both the simulated and empirical datasets with 20 taxa exhibited similar patterns. For a given number of informative sites, increasing the number of taxa was associated with greater difficulty in inferring the topology. Therefore, when comparing alignments with the same number of sequences, those containing more informative sites are more likely to yield better phylogenetic resolution.

**Figure 5.**
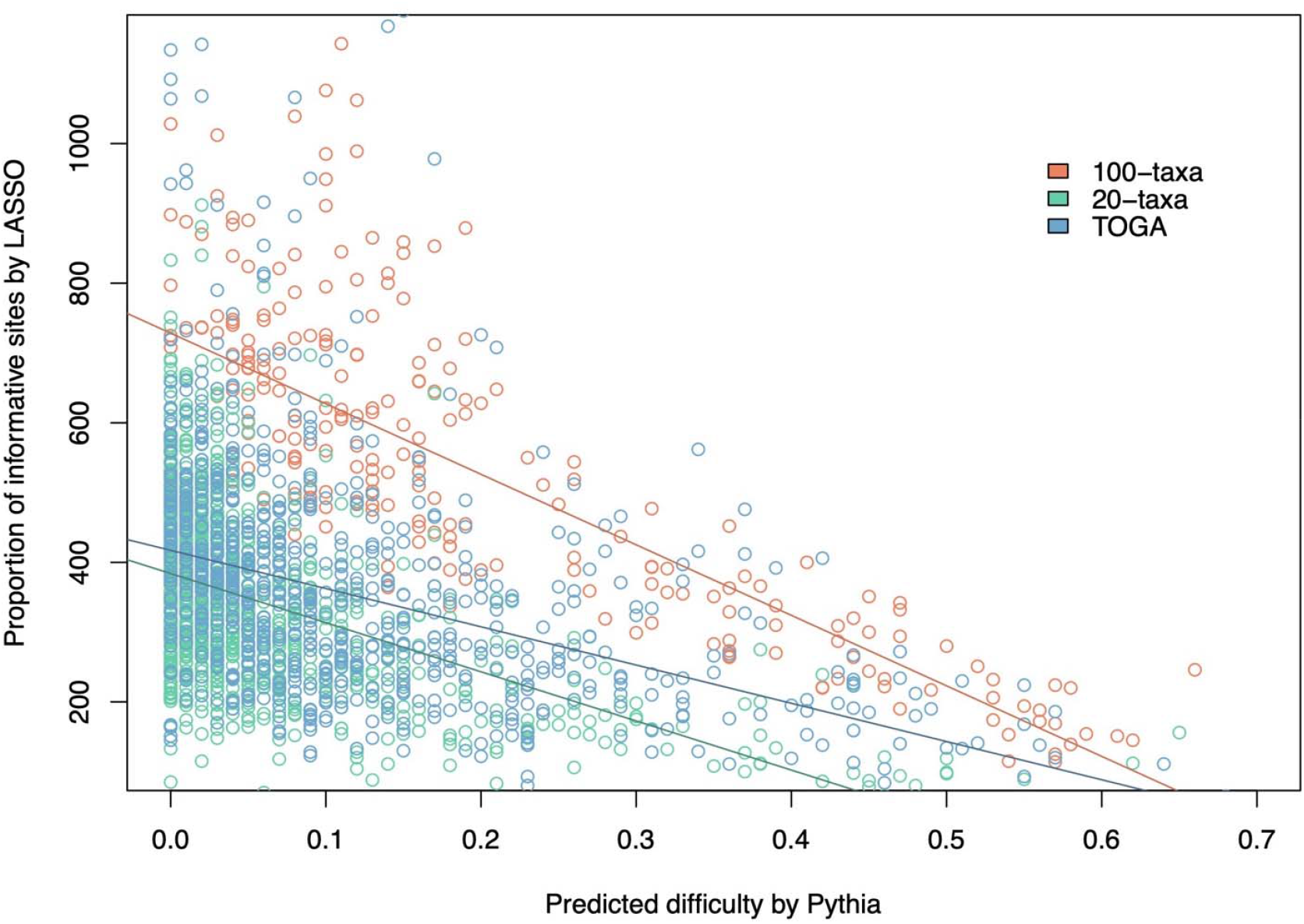
Inverse correlation between the proportion of informative sites (as identified by Lasso) and the predicted difficulty of phylogenetic inference, as measured by the Pythia model (Haag *et al*., 2022). Alignments with more informative sites consistently showed lower predicted difficulty across all datasets.

## Discussion

In this study, we introduced a sparse learning framework using the Lasso regressor to evaluate the phylogenetic information content of individual alignment sites. Our approach explicitly identifies the subset of sites that most strongly contribute to resolving the tree topology. Despite their significantly reduced length, alignments filtered to contain only Lasso-selected sites exhibited phylogenetic resolution close to that of the entire sequence. Our simulations also indicated that tree topologies inferred from Lasso-trimmed alignments were as close to the true trees as those inferred from the complete alignments. Using the Lasso-derived site-wise classification, we identified a simple Shannon entropy cut-off that approximated the results from the time-consuming Lasso procedure. Alignments filtered to retain only phylogenetically informative sites were, on average, one-fourth of the original length, potentially accelerating the computational time of phylogenomic pipelines. Although our empirical case study is mammalian, the mechanics of site-wise likelihood profiling and an information-theoretic proxy are lineage-agnostic.

Our findings confirm that phylogenetic signal is highly unevenly distributed among sites (Townsend et al., 2012). A small subset of sites exerts a disproportionate influence on the phylogenetic likelihood, a pattern also emphasized by Shen et al. (2017) and Walker et al. (2018). In this sense, our results differ from the conclusions of Tan et al. (2015), who reported that filtering alignments degrades the phylogenetic signal. We found that, on average, even removing more than 75% of alignment sites, the data retained information that could resolve the tree topology accurately (**Figure 2a**). Moreover, branch supports were close to those calculated for the complete alignment (**Figure 2b**).

We confirmed that gap-containing alignment sites carried significant phylogenetic information. In the mammalian dataset analyzed, 51.4% of sites had a gap frequency between 0.0 and 0.5, while 47.2% of Lasso-selected informative sites were gapless. Besides Shannon entropy, this finding could provide another practical criterion for site selection, i.e., gap frequency ≤ 0.5, and some studies have indeed adopted this criterion (Vanderpool *et al*., 2020; Stiller *et al*., 2024). However, applying this gap frequency threshold to mammalian alignments from TOGA would have retained 73.6% of the sites identified as uninformative by Lasso, because these sites are primarily conserved regions with a few autapomorphic substitutions that contribute little to resolving phylogenetic relationships.

Our conceptualization of phylogenetic information was primarily operational. Accordingly, our suggestion to define the phylogenetically effective alignment length as the subset of sites that significantly contribute to the tree likelihood is empirically motivated. This contrasts with the recent theoretical framework introduced by Seo et al. (Seo *et al*., 2022), who defined the effective sequence length as the length of a hypothetical gapless alignment that retains the same amount of phylogenetic information as the actual (gap-containing) alignment. By grounding their approach in Fisher information, they introduced a formal analysis of the partitioned impact of gaps and model misspecification on phylogenetic information by analyzing the curvature of the likelihood function. While their treatment differs from ours, they proposed a site-specific metric (s-ESL) for quantifying information content, which is comparable to our Lasso-based approach. In agreement with our results, Seo et al. (2022) also found that gaps do not necessarily obliterate phylogenetic information.

Because we provided a standard quantification of site-wise phylogenetic information, we can independently assess the performance of entropy-based metrics in phylogenomics. Our Lasso classification revealed that most high-entropy sites contributed significantly to the tree likelihood (**Figure 3a**). This finding suggests that these sites should not be dismissed as uninformative a priori (Ballesteros *et al*., 2022). For instance, sites with evenly distributed character states exhibit high entropy. While such patterns may appear noisy, they can carry meaningful phylogenetic signal when associated with the tree topology. Therefore, site entropy alone, when assessed independently of a tree, is not a reliable measure of phylogenetic noise. Entropy-based analyses should be interpreted in conjunction with phylogenetic likelihood to quantify site informativeness in multiple sequence alignments accurately.

Although our analysis has not addressed site-level saturation, Duchêne et al. (2022) have demonstrated that substitution saturation is a pervasive source of error in phylogenomic inference, even under well-fitted substitution models. They addressed this by developing an entropy-based test to flag and exclude entire loci with excessive saturation. In our study, while we did not perform locus-level saturation diagnostics, the number of informative sites provides a valuable proxy for locus quality. Its positive correlation with an independent predictor of phylogenetic utility, the Pythia model, supports this approach. When building phylogenomic pipelines, it may be beneficial to integrate locus-level saturation screening with site-wise, model-aware filtering strategies. Such a combined approach could enhance the accuracy and reliability of downstream phylogenetic inference.

From a phylogenomic design perspective, our findings underscore the importance of sequence length. We observed a positive correlation between the number of informative sites and alignment length, suggesting that longer loci increase the probability of accurately resolving gene tree topologies. This supports the idea that phylogenomic pipelines should prioritize assembling longer genes sampled from diverse genomic regions. However, while longer genomic segments tend to carry more phylogenetic information, they are also more prone to intragenic recombination. This increases the risk of combining sites with conflicting evolutionary histories, potentially complicating analyses under the multispecies coalescent (MSC) model (Gatesy and Springer, 2013). As such, it remains an open question whether datasets with longer genes are more likely to recover the correct species tree topology. Alternatively, recent MSC methods that infer species trees directly from site pattern frequencies, thereby bypassing gene tree estimation, may offer a more robust framework (Zhang *et al*., 2025).

In conclusion, our approach represents an advancement in quantifying and harnessing phylogenetic information at the site level. It offers a scalable, interpretable, and principled tool for alignment refinement, conflict diagnosis, and experimental design in phylogenomics, while also highlighting key avenues for further methodological development. Our implementation of the Lasso regressor is available at http://github.com/cschrago/LassoTrim.

## Acknowledgements

CGS is supported by grants 309165/2019-9, 409963/2023-2, and 302910/2025-5 from the Conselho Nacional de Desenvolvimento Científico e Tecnológico (CNPq).

