## Supplementary material for "Efficient Identification of Phylogenetically Informative Alignment Sites via Sparse Learning": Table S1

**Table S1.** Frequency at which Lasso and Lasso-based entropy (*H*) alignments outperformed or matched the performance of two trimming software, as measured by the RF distances between the true topology and the IQ-TREE2-inferred trees (*d*_Lasso/_*_H_* $\leq$ *d*_trimAl/BMGE_).

|  | trimAl | BMGE |
| --- | --- | --- |
| Lasso |  |  |
| 20-taxa | 0.62 | 0.6 |
| 100-taxa | 0.51 | 0.51 |
| *H* |  |  |
| 20-taxa | 0.72 | 0.72 |
| 100-taxa | 0.31 | 0.32 |
